# Genomic GC content drifts slowly downward in most bacterial genomes

**DOI:** 10.1101/2020.07.19.211128

**Authors:** Bert Ely

**Affiliations:** Department of Biological Sciences, University of South Carolina, Columbia, SC, 29203 USA

**Keywords:** genomic GC content, mutation accumulation, mutation bias, prokaryotes, Caulobacter

## Abstract

In every kingdom of life, GC->AT transitions occur more frequently than any other type of mutation due to the spontaneous deamination of cytidine. In eukaryotic genomes, this slow loss of GC base pairs is counteracted by biased gene conversion which increases genomic GC content as part of the recombination process. However, this type of biased gene conversion has not been observed in bacterial genomes so we hypothesized that GC->AT transitions cause a reduction of genomic GC content in prokaryotic genomes on an evolutionary time scale. To test this hypothesis, we used a phylogenetic approach to analyze triplets of closely related genomes representing a wide range of the bacterial kingdom. The resulting data indicate that genomic GC content is slowly declining in bacterial genomes where GC base pairs comprise 40% or more of the total genome. In contrast, genomes containing less than 40% GC base pairs have fewer opportunities for GC->AT transitions to occur so genomic GC content is relatively stable or actually increasing at a slow rate. It should be noted that this observed change in genomic GC content is the net change in shared parts of the genome and does not apply to parts of the genome that have been lost or acquired since the genomes being compared shared common ancestor. However, a more detailed analysis of two *Caulobacter* genomes revealed that the acquisition of mobile elements by the two genomes actually reduced the total genome content as well.

## Introduction

Bacterial genomic GC content varies from less than 15% to more than 75% [1]. Furthermore, a wide range of genomic GC content can be observed among the genera within most bacterial phyla. However, genomic GC content usually varies in a narrow range within a single bacterial genus indicating that genomic GC content is relatively constant over evolutionary time scales [1]. In contrast, mutations that occur in bacterial genomes accumulate over time causing changes that have led to the diversification of individual species within a genus. Genomic GC content is maintained by mutational forces that are balanced by mutations occurring during DNA replication and DNA repair [2]. For example, Wu et al. [3] proposed that the *polC* and *dnaE2* genes have a great impact on genomic GC content. However, mutations that convert GC base pairs to AT base pairs (GC->AT) occur more frequently than any other type of mutation, primarily due to spontaneous cytidine deamination [4, 5]. Therefore, cytidine deamination could be a driving force that could lead to a reduction of genomic GC content over evolutionary time periods. Several hypotheses have been proposed to explain how bacteria are able to maintain genomes with high percentages of GC base pairs and counteract the cytidine deamination mutation pressure [4-8]. For example, the impact of the conversion of GC base pairs to AT base pairs could be counteracted by the process of horizontal gene transfer (HGT) to import GC-rich DNA segments and maintain the overall GC content of the genome. However, this type of HGT would result in patches of imported DNA with higher GC content while the rest of the genome had a continually decreasing GC content. In addition, there would have to be some sort of selection for the import of GC-rich segments into the bacterial genome. In contrast, several analyses have shown that most imported genomic regions obtained via HGT in a variety of bacteria come from closely-related genomes with similar GC content [9-13]. Also, careful analyses of closely related *Caulobacter vibrioides* genomes failed to reveal imported regions with higher than average GC content [14]. Therefore, HGT is not likely to be the primary way that high average GC content is maintained in bacterial genomes.

An alternative hypothesis, biased gene conversion, suggests that AT base pairs are converted to GC base pairs in mismatched regions of DNA that are generated during recombination events. Since this phenomenon has been well established for eukaryotic genomes [15-18], it has been considered a reasonable hypothesis, and some data have been presented that are consistent with the idea that biased gene conversion has occurred in bacterial genomes [19]. If biased gene conversion did occur in bacteria, then recombination events could help maintain a constant genomic GC content by causing AT-> GC transitions in sufficient quantity to compensate for the excess GC->AT transitions. However, biased gene conversion could only occur during relatively rare recombination events [14] while eukaryotic recombination occurs multiple times during every round of meiosis. Furthermore, we have shown that biased gene conversion did not occur during more than 100 distinct recombination events analyzed in the high GC genomes of *C. vibrioides* [20]. Thus, biased gene conversion does not appear to occur during recombination events in the high GC genomes of this bacterial species and recombination does not occur frequently enough to counteract the observed rates of cytidine deamination in most bacterial species.

If recombination and HGT do not have a significant impact genomic GC content, then how is high GC content maintained in bacterial genomes? One proposed explanation is that there is some type of selection for higher genomic GC content [4-8]. However, although many possibilities have been ruled out, no one has identified a selective mechanism that would increase genomic GC content. It is possible that repair processes are sufficiently biased to counteract the ongoing cytidine deamination so the GC->AT pairs resulting from deamination are precisely balanced by AT->GC conversions occurring during repair. Alternatively, we hypothesized that bacterial genomic GC content actually is increasing or decreasing due to the GC->AT mutation pressure from cytidine deamination. To test this hypothesis, we conducted a series of phylogenetic analyses of polymorphisms that have accumulated over millions of generations in a wide variety of prokaryotic genera. These analyses demonstrated that prokaryotic genomes with high or average levels of GC content are slowly evolving to lower levels of GC content, but that the pace of this change is extremely slow so that genomic GC content appears to be constant.

## Methods

### Genome comparisons

A wide range of representative prokaryotic species were chosen based on the availability of three closely-related and complete genome sequences in the GenBank database. If available, we chose sets of three closely-related genomic sequences from two genera from each bacterial phylum as well as several from each class of the *Proteobacteria* (Table 1). Each set of three related genomes was aligned with progressiveMauve using standard settings, and the positions of all single nucleotide polymorphisms (SNPs) were exported as a single file that showed which base was present in each of the three genomes at each of the variable locations [21]. The more distantly related genome was identified from the phylogenetic trees that are present for each species in the NCBI database. Using this phylogenetic approach, the SNPs that differed between the two more closely related genomes were compared, and the SNP that matched the SNP in the third, more distal genome was considered the ancestral allele. Positions that contained three different alleles were omitted from the analysis since the ancestral state could not be determined. Also, genome comparisons that generated fewer than 1000 SNPs with defined ancestral alleles were omitted from further analyses due to the small sample size. Each combination of the ancestral and derived alleles was tallied over the whole genome, and the G->A, G->T, C->A, and C->T changes were summed to generate total GC->AT transitions. Likewise, the A->G, A->C, T->G, and T->C changes were summed to generate the total AT->GC transitions. The remaining SNPs, A->T, T->A, G->C, and C->G resulted in no change in GC content and were not considered further. To calculate the net change in genomic GC content, the number of AT->GC SNPs was subtracted from the number of GC->AT SNPs to determine the net loss of GC base pairs in the genome. (Therefore, a negative number would indicate an increase in genomic GC content.) The net change in the number of GC base pairs was divided by the size of the bacterial genome to determine the percentage change in the genomic GC content since the time when the two bacterial strains shared a common ancestor. Also, the net change of genomic GC content was divided by the total number of SNPs analyzed to facilitate comparisons of the change in GC content among the analyzed species to compensate for differences in time since the most recent common ancestor.

**Table 1.** Prokaryotic species used in this study.

| **Phylum** | **Species** | **Proteobacteria** | **Species** |
| --- | --- | --- | --- |
| *Actinobacteria* | *Corynebacterium diphtheriae* | *Alphaproteobacteria* | *Caulobacter vibrioides* |
|  | *Cutibacterium acnes* |  | *Candidatus Sulcia muelleri* |
| *Bacteroidetes* | *Flavobacterium psychrophilum* |  | *Ehrlichia chaffeensis* |
|  | *Rhodothermus marinus* |  | *Ochrobactrum anthropi* |
| *Chlamydiae* | *Chlamydia pecorum* | *Betaproteobacteria* | *Polynucleobacter asymbioticus* |
|  | *Chlamydia trachomatis* | *Gammaproteobacteria* | *Haemophilus influenza* |
|  | *Chlamydia psittaci* |  | *Pseudomonas putida* |
|  | *Chlamydia suis* |  | *Escherichia coli* |
| *Cyanobacteria* | *Prochlorococcus marinus* | *Deltaproteobacteria* | *Desulfovibrio vulgaris* |
|  | *Synechococcus sp.* |  |  |
| *Deinococcus-Thermus* | *Thermus thermophilus* |  |  |
| *Firmicutes* | *Bacillus cereus* |  |  |
|  | *Bacillus subtilis* |  |  |
| *Fusobacteria* | *Fusobacterium necrophorum* |  |  |
|  | *Fusobacterium hwasookii* |  |  |
| *Spirochaetes* | *Brachyspira pilosicoli* |  |  |
|  | *Brachyspira hyodysenteriae* |  |  |
| *Thermotogae* | *Thermotoga* |  |  |
| *Verrucomicrobia* | *Akkermansia muciniphila* |  |  |
| *Archaea* | *Methanococcus maripaludis* |  |  |
|  | *Thermococcus sp.* |  |  |

## Results and Discussion

Since our laboratory specializes in C*aulobacter* genetics, we first analyzed closely related *C. vibrioides* genomes that had been recently sequenced [14]. To determine if genomic GC content has remained constant, we identified 73,859 SNPs by comparing the CB2 and NA1000 genomes and then identified the ancestral allele of 46,043 of those SNPs by comparison of the SNPs to the nucleotides at the corresponding positions in the more distantly-related CB13 genome [14]. The regions corresponding to the positions of the remaining SNPs were not present in the CB13 genome so the ancestral allele could not be identified.

We found that GC->AT changes were the most common (49% of the 46,043 SNPs) while AT->GC SNPs occurred at a frequency of only 21%. Thus, in each strain, there has been a net loss of GC base pairs that would have caused a reduction in the genomic GC content of about 0.15% during the time period since their most recent common ancestor (Table 2), which is estimated to have existed 25 million generations ago [14]. To ensure that these calculations were not influenced by the choice of the reference genome, we repeated the analysis using the more distantly-related *C. segnis* TK0057 genome [14, 22] as the reference genome and found a 0.2% reduction of the genomic GC content since CB2 and NA1000 shared a common ancestor (data not shown). Although the magnitude of the genomic GC content reduction is similar in the two analyses, the first estimate is probably more accurate since we were able to determine the ancestral state of nearly 50% more SNPs when the more closely-related CB13 genome was used as the reference. It is important to note that this rate of decrease in genomic GC content reflects the SNPs that have accumulated over time throughout the entire genome. These SNPs are the net result of the mutations that occur over time with selection for more favorable genetic content and genetic drift that is independent of selection.

**Table 2.** Loss of GC content in *Alphaproteobacteria*.

| Phyla and Bacteria | Net Loss of GC Content | Total Mutations | Genome Size (Mb) | Genomic %GC Content | Percent Change in GC | SNP % of Genome | Loss of GC/SNPs |
| --- | --- | --- | --- | --- | --- | --- | --- |
| ***Caulobacter vibrioides*** |  |  |  |  |  |  |  |
| *C. vibrioides* NA1000 | 6420 | 23250 | 4.0 | 67.7 | 0.16 | 0.58 | 0.276 |
| *C. vibrioides* CB2 | 6310 | 22793 | 4.1 | 67.2 | 0.15 | 0.56 | 0.277 |
| *C. vibrioides* CB13 |  |  | 4.1 | 67.1 |  |  |  |
| ***Escherichia coli*** |  |  |  |  |  |  |  |
| *E. coli* P12b | 1934 | 12722 | 4.9 | 50.9 | 0.04 | 0.26 | 0.152 |
| *E. coli* K-12 | 1734 | 10894 | 4.6 | 50.8 | 0.04 | 0.24 | 0.159 |
| *E. coli* 142 |  |  | 5.7 | 50.5 |  |  |  |
| *E. coli* 142 | 125 | 328 | 5.7 | 50.5 | 0.002 | 0.06 | 0.03 |
| *E. coli* 0157 644-PT8 | 119 | 247 | 5.8 | 50.5 | 0.002 | 0.04 | 0.05 |
| *E. coli* K-12 |  |  | 4.6 | 50.8 |  |  |  |

A limitation of this approach is that only the SNPs that are also present in the strain that serves as the reference can be analyzed. However, as indicated above, we were able to analyze 62% of the total number of SNPs present in the comparison of the CB2 and NA1000 genomes. If we assume that the SNPs that we analyzed are representative of all the SNPs identified in the two genomes, then our estimate of the loss of GC content might be only 62% of the actual loss of GC content. Also, both the CB2 and NA1000 genomes contain regions that are not present in the other genome. However, these regions represent only 8% of the two genomes, and more than half of these differences are due to eight mobile elements that are unique to one of the two genomes (four in each genome) and are not present in the CB13 genome. These mobile elements have a slightly lower GC content than the genome average and together would reduce the total genomic GC content by an additional 0.1%. Taken together, these data indicate that the GC content of the *C. vibrioides* NA1000 and CB2 genomes has been slowly drifting downward as a result of the insertion of mobile elements and the accumulation of GC->AT transitions. The net result of all of these changes is that the common ancestor of the *C. vibrioides* NA1000 and CB2 wild type strains lived about 25 million generations ago [11] and its estimated genomic GC content was approximately 0.3% percent higher than the current genomic GC content.

These results contrast with previous results that showed that AT->GC mutations were the most common types of mutations in *C. vibrioides* [23, 24]. However, in these previous studies, the number of mutations was both small and limited by selection for a particular phenotype (rifampicin resistance or inactivation of the XylR repressor). Thus, the mutations selected under the conditions used in these two laboratory studies do not appear to be representative of the mutations that accumulate throughout the entire genome over evolutionary time periods.

To determine if slow losses of GC content also had occurred in *Escherichia coli*, we performed similar analyses using the genomes of four *E. coli* strains (Table 2). In the first comparison, when the two closely related strains differed by more than 10,000 SNPs, the genomic GC content decreased by approximately 0.04%. In a second comparison, when the closely related strains differed by only 575 total SNPs, the decrease in genomic GC content was only 0.002%. Thus, the decrease in the genomic GC content is roughly proportional to the number of SNPs examined in the two situations.

Based on the results of the *C. crescentus* and *E. coli* analyses, we hypothesized that loss of genomic GC content might be widespread among prokaryotic genera. Therefore, we examined genomes from 28 bacterial species representing 11 different phyla as well as those of two archaeal genera (Tables 1 and S1). Two simultaneous trends were observed: 1) Greater loss of GC content occurred when the genomic GC content was higher (Fig. 1); and 2) greater loss of GC content occurred when the genetic distance between the two genomes was greater (represented by the total number of SNPs observed between the two genomes being compared) (Fig. 2). Both of these trends are consistent with expectations. A larger number of SNPs indicates that a longer time had elapsed since the two strains shared a common ancestor so that there would have been more time for an excess of GC ->AT transitions to accumulate. For example, the six comparisons with the highest numbers of SNPs as a percentage of the genome also were the farthest from the Y axis indicating that they had experienced the greatest changes in GC content (Fig. 2). Similarly, a higher GC content would provide more opportunities for cytidine deamination to occur. Thus, for genomes with a GC content greater than 40%, there was a trend towards increasing loss of GC content after correcting for the total number of mutations relative to the size of the genome (Fig. 1). Based on these results, we conclude that a slow decrease in genomic GC content is a common phenomenon for prokaryotic genomes with a genomic GC content that is greater than 40%.

**Fig. 1.**
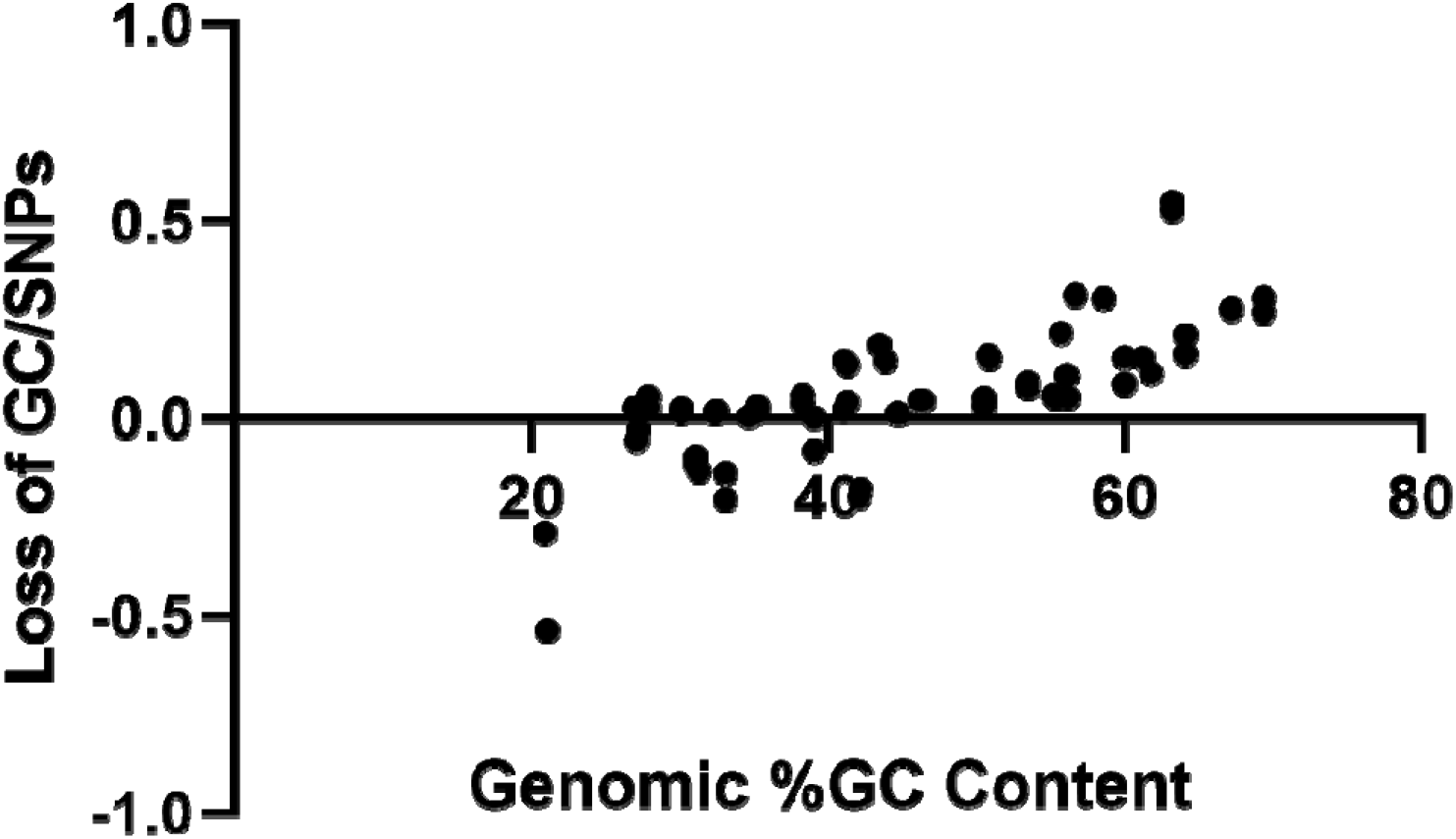
Net change in genomic GC content divided by the total number of SNPs analyzed versus the genomic GC content.

**Fig. 2.**
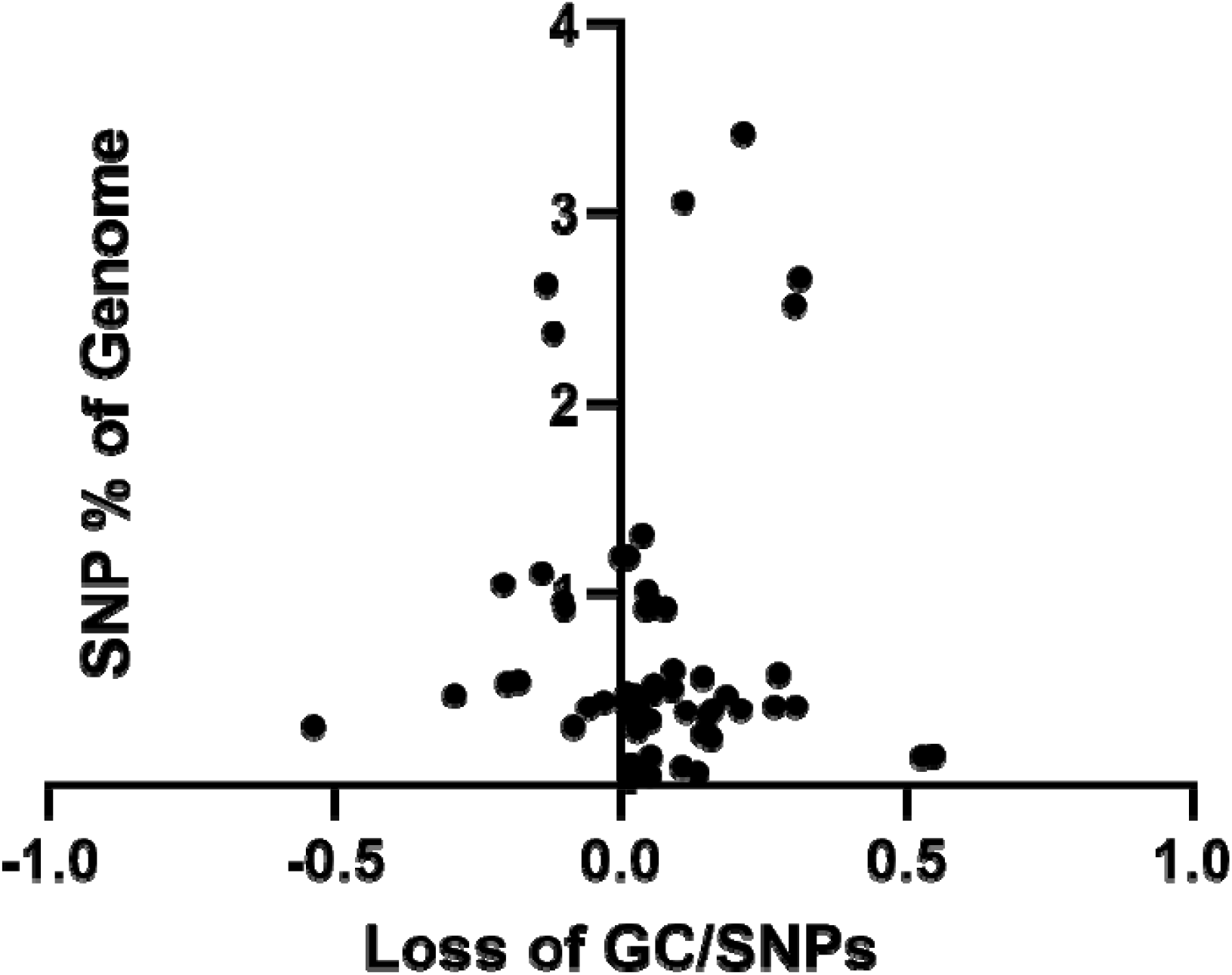
Change in genomic GC content (X axis) versus the number of SNPs analyzed relative to the size of the genome.

When we analyzed prokaryotic genomes with a genomic GC content that was less than 40%, five genera had a surplus of AT->GC transitions resulting an increase in genomic GC content, and five had a net decrease in genomic GC content (Table 1). In addition, one strain in the *Fusobacterium* comparison had accumulated a slight increase in genomic GC content while the other had accumulated a slight decrease in genomic GC content. However, in each of these 11 genera, the number of observed GC->AT transitions was higher per GC base pair than the corresponding number of AT->GC transitions per AT base pair. Thus, these data provide further evidence that the conversion of GC base pairs to AT base pairs occurs at a higher rate in all prokaryotic genomes regardless of GC content. Cytidine deamination is probably the most important factor contributing to the higher rate of GC->AT transitions, but the impact of these transitions on genomic GC content is dependent on the percentage of GC base pairs present in the genome. Thus, if the genomic GC content is low enough, the AT->GC mutations would outnumber GC->AT mutations due to the much smaller number of GC base pairs that are present in the genome.

If high genomic GC content is slowly reduced due to GC->AT mutation pressure, then how did high GC bacterial genomes arise? Our previous analysis of bacterial phyla demonstrated that genomic GC content was fairly constant in most genera, but that wide variations in genomic GC content are present among genera within phyla [1]. Thus, during the evolution of bacterial genera within phyla, there may have been a period of time when repair systems had a wide range of AT->GC conversions and conditions favored divergent genomic evolution patterns. As a result, the mutation pressure and selection that occurred under those conditions might have resulted in the differentiation of genomic GC content among genera within the same phylum.

## Acknowledgements

I thank Dakota Lee for helping with many of the genome comparisons.

