## Supplemental Table 1 for "Genomic GC content drifts slowly downward in most bacterial genomes"

**Supplementary Table 1.** Phylogenetic analyses of SNPs present in closely related prokaryotic genomes.

| Phyla and Bacteria | Net Loss of GC Content | Total Mutations | Genome Size (Mb) | Genomic %GC Content | Percent Change in GC | SNP % of Genom[illegible] |
|---|---|---|---|---|---|---|
| ***Proteobacteria*** | | | | | | |
| ***Alphaproteobacteria*** | | | | | | |
| ***Caulobacter vibrioides*** | | | | | | |
| *C. vibrioides* NA1000 | 6420 | 23250 | 4.0 | 67.7 | 0.16 | 0.5[illegible] |
| *C. vibrioides* CB2 | 6310 | 22793 | 4.1 | 67.2 | 0.15 | 0.5[illegible] |
| *C. vibrioides* CB13 | | | 4.1 | 67.1 | | |
| ***Candidatus Sulcia muelleri*** | | | | | | |
| *C. Sulcia muelleri* CARI | -439 | 817 | 0.28 | 21.1 | -0.16 | 0.3[illegible] |
| *C. Sulcia muelleri* PSPU | -392 | 1346 | 0.29 | 20.9 | -0.14 | 0.4[illegible] |
| *C. Sulcia muelleri* DMIN | | | 0.24 | 22.5 | | |
| ***Ehrlichia chaffeensis*** | | | | | | |
| *E. chaffeensis* Liberty | 13 | 517 | 1.18 | 30.1 | 0.00 | 0.0[illegible] |
| *E. chaffeensis* Jax | 7 | 449 | 1.18 | 30.1 | 0.00 | 0.0[illegible] |
| *E. chaffeensis* Heartland | | | 1.17 | 30.1 | | |
| ***Ochrobactrum anthropi*** | | | | | | |
| *O. anthropi* ATCC 49188 | 367 | 7092 | 5.2 | 56.2 | 0.01 | 0.1[illegible] |
| *O. anthropi* OAB | 475 | 4502 | 4.9 | 56.1 | 0.01 | 0.0[illegible] |
| *O. anthropi* T16R-87 | | | 4.7 | 56.0 | | |
| ***Betaproteobacteria*** | | | | | | |
| ***Polynucleobacter asymbioticus*** | | | | | | |
| *P. asymbioticus* Tro8F10W22 | 111 | 10736 | 2.22 | 44.8 | 0.00 | 0.4[illegible] |
| *P. asymbioticus* MWH-RechtKolB | 380 | 28301 | 2.36 | 44.7 | 0.02 | 1.2[illegible] |
| *P. asymbioticus* QLW-P1DMWA-1 | | | 2.16 | 44.8 | | |
| ***Gammaproteobacteria*** | | | | | | |
| ***Haemophilus influenza*** | | | | | | |
| *H. influenza* KW20 | 992 | 17101 | 1.83 | 38.3 | 0.05 | 0.9[illegible] |
| *H. influenza* 2019 | 950 | 25902 | 1.97 | 38.3 | 0.05 | 1.3[illegible] |
| *H. influenza* F3031 | | | 1.99 | 38.2 | | |
| ***Pseudomonas putida*** | | | | | | |
| *P. putida* F1 | 3269 | 21361 | 6.0 | 61.2 | 0.05 | 0.3[illegible] |
| *P. putida* ND6 | 2620 | 22956 | 6.1 | 61.8 | 0.04 | 0.3[illegible] |
| *P. putida* DOT-T1E | | | 6.3 | 61.4 | | |
| ***Escherichia coli*** | | | | | | |
| *E. coli* K-12 | 5533 | 28529 | 4.6 | 50.8 | 0.12 | 0.6[illegible] |
| *E. coli* 1428 | 6122 | 74473 | 5.3 | 50.6 | 0.12 | 1.4[illegible] |
| *E. coli* O104H4 | | | 5.3 | 50.8 | | |

| | | | | | | |
|---|---|---|---|---|---|---|
| ***Deltaproteobacteria*** | | | | | | |
| ***Desulfovibrio vulgaris*** | | | | | | |
| *D. vulgaris* DP4 | 2965 | 5425 | 3.7 | 63.2 | 0.08 | [illegible] |
| *D. vulgaris* RCH1 | 2850 | 5407 | 3.7 | 63.2 | 0.08 | [illegible] |
| *D. vulgaris* Myazaki F | | | 4.0 | 67.1 | | |
| ***Actinobacteria*** | | | | | | |
| ***Corynebacterium diphtheriae*** | | | | | | |
| *C. diphtheriae* C7 (beta) | 1769 | 22991 | 2.50 | 53.5 | 0.07 | [illegible] |
| *C. diphtheriae* NCTC7838 | 1329 | 14732 | 2.44 | 53.5 | 0.05 | [illegible] |
| *C. diphtheriae* HC03 | | | 2.44 | 53.5 | | |
| ***Cutibacterium acnes (Propionibacterium)*** | | | | | | |
| *C. acnes* KPA171202 | 999 | 6478 | 2.56 | 60.0 | 0.04 | [illegible] |
| *C. acnes* TypeIA2 P.acn31 | 1091 | 12435 | 2.50 | 60.0 | 0.04 | [illegible] |
| *C. acnes subsp. defendens* ATCC 11828 | | | 2.49 | 60.0 | | |
| ***Bacteroidetes*** | | | | | | |
| ***Flavobacterium psychrophilum*** | | | | | | |
| *F. psychrophilum* JIP02/86 | 63 | 2107 | 2.90 | 32.5 | 0.002 | [illegible] |
| *F. psychrophilum* CSF259-43 | 13 | 565 | 2.90 | 32.5 | 0.000 | [illegible] |
| *F. psychrophilum* FPG101 | | | 2.80 | 32.5 | | |
| *F. psychrophilum* FPG101 | 57 | 2941 | 2.80 | 32.5 | 0.002 | [illegible] |
| *F. psychrophilum* JIP02/86 | 10 | 476 | 2.90 | 32.5 | 0.000 | [illegible] |
| *F. psychrophilum* V4-24 | | | 2.70 | 32.4 | | |
| ***Rhodothermus marinus*** | | | | | | |
| *R. marinus* AA2-13 | 2190 | 13490 | 3.44 | 64.1 | 0.06 | [illegible] |
| *R. marinus* AA3-38 | 2796 | 13242 | 3.43 | 64.1 | 0.08 | [illegible] |
| *R. marinus* SG0.5JP17-172 | | | 3.33 | 64.3 | | |
| ***Chlamydiae*** | | | | | | |
| ***Chlamydia pecorum*** | | | | | | |
| *C. pecorum* E58 | 421 | 2912 | 1.11 | 41.1 | 0.04 | [illegible] |
| *C. pecorum* P787 | 82 | 4170 | 1.11 | 41.1 | 0.01 | [illegible] |
| *C. pecorum* PV30563 | | | 1.10 | 41.1 | | |
| ***Chlamydia trachomatis*** | | | | | | |
| *C. trachomatis SQ29* | 90 | 668 | 1.05 | 41.3 | 0.01 | [illegible] |
| *C. trachomatis E-DK-20* | 33 | 826 | 1.06 | 41.3 | 0.00 | [illegible] |
| *C. trachomatis D-LC* | | | 1.05 | 41.3 | | |
| ***Chlamydia psittaci*** | | | | | | |
| *C. psittaci* 6BC | -4 | 14128 | 1.18 | 39.1 | 0.00 | [illegible] |
| *C. psittaci* GR9 | -283 | 3434 | 1.15 | 39.1 | -0.02 | [illegible] |
| *C. psittaci* WSRTE30 | | | 1.15 | 39.0 | | |
| ***Chlamydia suis*** | | | | | | |
| *C. suis* 3-25b | -1144 | 5737 | 1.08 | 42.1 | -0.11 | [illegible] |

| | | | | | | |
|---|---|---|---|---|---|---|
| *C. suis* 5-27b | -1076 | 5964 | 1.10 | 42.2 | -0.10 | 0.5[illegible] |
| *C. suis* 9-1b | | | 1.10 | 42.2 | | |
| ***Cyanobacteria*** | | | | | | |
| ***Prochlorococcus marinus*** | | | | | | |
| *P. marinus* MIT9202 | -1524 | 15623 | 1.69 | 31.1 | -0.09 | 0.9[illegible] |
| *P. marinus* MIT9215 | -1713 | 16725 | 1.74 | 31.1 | -0.10 | 0.9[illegible] |
| *P. marinus* MIT604 | | | 1.78 | 31.1 | | |
| *P. marinus* MIT9301 | -5684 | 43153 | 1.64 | 31.3 | -0.35 | 2.6[illegible] |
| *P. marinus* AS9601 | -4672 | 39623 | 1.67 | 31.1 | -0.28 | 2.3[illegible] |
| *P. marinus* MIT9215 | | | 1.74 | 31.2 | | |
| ***Synechococcus sp.*** | | | | | | |
| *Synechococcus sp.* CC9902 | 18302 | 58502 | 2.20 | 56.7 | 0.83 | 2.6[illegible] |
| *Synechococcus sp.* BL107 | 17623 | 58007 | 2.30 | 58.6 | 0.77 | 2.5[illegible] |
| *Synechococcus sp.* CC9605 | | | 2.50 | 59.2 | | |
| ***Deinococcus-Thermus*** | | | | | | |
| ***Thermus thermophilus*** | | | | | | |
| *T. thermophilus* HB27 | 2356 | 8747 | 2.13 | 69.4 | 0.11 | 0.4[illegible] |
| *T. thermophilus* HC11 | 2754 | 8994 | 2.17 | 69.4 | 0.13 | 0.4[illegible] |
| *T. thermophilus* TMY | | | 2.14 | 69.0 | | |
| ***Firmicutes*** | | | | | | |
| ***Bacillus cereus*** | | | | | | |
| *B. cereus* 14579 | 444 | 23384 | 5.4 | 35.3 | 0.01 | 0.4[illegible] |
| *B. cereus* FORC 013 | 726 | 23562 | 5.7 | 35.2 | 0.01 | 0.4[illegible] |
| *B. cereus* HN001 | | | 5.4 | 35.3 | | |
| ***Bacillus subtilis*** | | | | | | |
| *B. subtilis* 168 | 3561 | 19171 | 4.2 | 43.5 | 0.08 | 0.4[illegible] |
| *B. subtilis* BSP1 | 3244 | 22344 | 4.0 | 43.9 | 0.08 | 0.5[illegible] |
| *B. subtilis* RO-NN-1 | | | 4.0 | 43.9 | | |
| ***Fusobacteria*** | | | | | | |
| ***Fusobacterium necrophorum*** | | | | | | |
| *F. necrophorum subsp. funduliforme* F1291 | 270 | 9287 | 2.14 | 35.3 | 0.01 | 0.4[illegible] |
| *F. necrophorum subsp. funduliforme* 1_1_36S | 47 | 9837 | 2.29 | 34.7 | 0.00 | 0.4[illegible] |
| *F. necrophorum* FDAARGOS_565 | | | 2.68 | 34.0 | | |
| ***Fusobacterium hwasookii*** | | | | | | |
| *F. hwasookii* ChDC F206 | -301 | 10511 | 2.43 | 27.2 | -0.01 | 0.4[illegible] |
| *F. hwasookii* ChDC F300 | -557 | 10015 | 2.53 | 27.1 | -0.02 | 0.4[illegible] |
| *F. hwasookii* ChDC F174 | | | 2.53 | 27.1 | | |
| ***Spirochaetes*** | | | | | | |
| ***Brachyspira pilosicoli*** | | | | | | |
| *B. pilosicoli* 951000 | 316 | 12273 | 2.59 | 27.9 | 0.01 | 0.4[illegible] |
| *B .pilosicoli* P43678 | 659 | 12191 | 2.56 | 27.9 | 0.03 | 0.4[illegible] |

| | | | | | | |
|---|---|---|---|---|---|---|
| *B. pilosicoli* B2904 | | | 2.77 | 27.8 | | |
| ***Brachyspira hyodysenteriae*** | | | | | | |
| *B. hyodysenteriae* BH718 | 233 | 8345 | 3.0 | 27.0 | 0.01 | 0.2 |
| *B. hyodysenteriae* WA1 | 251 | 10018 | 3.1 | 27.1 | 0.01 | 0.3 |
| *B. hyodysenteriae* B78 | | | 3.1 | 27.1 | | |
| ***Thermotogae*** | | | | | | |
| ***Thermotoga*** | | | | | | |
| *Thermotoga sp.* Cell2 | 806 | 17757 | 1.75 | 46.4 | 0.05 | 1.0 |
| *Thermotoga sp.* RQ2 | 766 | 17318 | 1.88 | 46.2 | 0.04 | 0.9 |
| *Thermotoga sp.* RQ7 | | | 1.85 | 47.1 | | |
| ***Verrucomicrobia*** | | | | | | |
| ***Akkermansia muciniphila*** | | | | | | |
| *A. muciniphila* EB-AMDK-10 | 821 | 14282 | 2.76 | 55.2 | 0.03 | 0.5 |
| *A. muciniphila* EB-AMDK-8 | 469 | 9253 | 2.82 | 55.4 | 0.02 | 0.3 |
| *A. muciniphila* MGYG-HGUT-02454 | | | 2.76 | 55.2 | | |
| ***Archaea*** | | | | | | |
| ***Methanococcus maripaludis*** | | | | | | |
| *M. maripaludis* S2 | -3569 | 17360 | 1.66 | 33.1 | -0.22 | 1.0 |
| *M. maripaludis* X1 | -2696 | 19426 | 1.75 | 33.1 | -0.15 | 1.1 |
| *M. maripaludis* C7 | | | 1.54 | 33.3 | | |
| ***Thermococcus sp.*** | | | | | | |
| *Thermococcus sp.* 5-4 | 13599 | 63274 | 1.85 | 55.7 | 0.74 | 3.4 |
| *Thermococcus sp.* 4457 | 6667 | 61433 | 2.01 | 56.1 | 0.33 | 3.0 |
| *Thermococcus sp.* AM4 | | | 2.09 | 54.8 | | |
